# Genome-wide association study of TP53 R249S mutation in hepatocellular carcinoma with aflatoxin B1 exposure and hepatitis B virus infection in Guangxi

**DOI:** 10.1101/2020.08.03.235135

**Authors:** Chuangye Han, Tingdong Yu, Wei Qin, Xiwen Liao, Jianlu Huang, Zhengtao Liu, Long Yu, Xiaoguang Liu, Zhiwei Chen, Chengkun Yang, Xiangkun Wang, Shutian Mo, Guangzhi Zhu, Hao Su, Zengnan Mo, Tao Peng

## Abstract

**Background/Aims:** Dietary aflatoxin B1 (AFB1) exposure, which induces DNA damage and codon 249 mutation of the TP53 gene, is one of the major risk factors for hepatocellular carcinoma (HCC). Hepatitis B virus (HBV) infection and AFB1 exert synergistic effects to promote carcinogenesis and TP53 R249S mutation in HCC.

**Methods:** A genome-wide association study (GWAS) was conducted on 485 cases of HCC with chronic HBV infection, followed by a two-stage replication study on 270 cases with chronic HBV infection. Susceptibility variants for the TP53 R249S mutation in HCC were identified based on both GWAS and replication analysis. The associations of identified variants with expression levels of their located genes were validated in 20 paired independent samples.

**Results:** Our results showed that TP53 R249S was significantly associated with ADAMTS18 rs9930984 (adjusted *P* = 4.84×10^−6^), WDR49 rs75218075 (adjusted *P* = 7.36 × 10^−5^) and SLC8A3 rs8022091 (adjusted *P* = 0.042). Additionally, ADAMTS18 mRNA expression was significantly higher in HCC tissue, compared with paired non-tumor tissue (*P* = 0.041) and patients carrying the TT genotype at rs9930984 showed lower ADAMTS18 expression in non-tumor tissue, compared with those carrying the GT genotype (*P* = 0.0028).

**Conclusions:** TP53 expression is significantly associated with R249S mutation in HCC. Our collective results suggest that rs9930984, rs75218075 and rs8022091 are associated with susceptibility to the R249S mutation in cases of HCC exposed to AFB1 and HBV infection.

## Introduction

Primary liver cancer is one of the most prevalent cancer type with over 841,000 new cases and causing approximately 782,000 cancer-related deaths worldwide in 2018. HCC represents 75%–85% of the histologic types of total liver cancer.(Bray *et al*. 2018) The main risk factors are chronic infection with HBV or hepatitis C virus, aflatoxin contamination of foodstuffs, heavy alcohol intake, obesity, smoking and type 2 diabetes.(El-Serag 2011; Mittal and El-Serag 2013) Chronic HBV infection is associated with 53% HCC worldwide(Perz *et al*. 2006) and a higher proportion (75%) of cases in Asia.(Huy and Abe 2004) AFB1, the major carcinogenic form of aflatoxin, has been classified as a group I carcinogen in humans by the International Agency for Research on Cancer (IARC)(Kujawa 1994) and a major risk factor for HCC. In China, 65.9% and 58.4% liver cancer-related deaths in men and women, respectively, were due to HBV infection, and an estimated 25.0% deaths attributable to aflatoxin exposure in both sexes in 2013.(Fan *et al*. 2013)

TP53, a tumor suppressor, responds to diverse cellular stress signals and regulates expression of target genes to induce cell cycle arrest, apoptosis, senescence, DNA repair or changes in metabolism. TP53 is the most commonly mutated gene in human malignancies.(Kandoth *et al*. 2013) TP53 mutations are reported to be associated with prognosis of various cancers and validated as biomarkers of environmental carcinogens.(Robles and Harris 2010) In particular, the R249S mutation in exon 7 of TP53 is a recognized molecular fingerprint of aflatoxin exposure(Smela *et al*. 2001; Liu *et al*. 2008; Forner *et al*. 2012; Qi *et al*. 2015) associated with poor prognosis of patients with HCC.(Woo *et al*. 2011) Previous reports suggest that HBV and AFB1 exposure have synergistic carcinogenic effects and increase the likelihood of TP53 R249S in HCC.(Kew 2003; Liu *et al*. 2008; Forner *et al*. 2012) Owing to high dietary exposure to AFB1 and HBV infection rates, the incidence of HCC in Guangxi is higher than the national level in China, with a frequency of TP53 R249S mutation of up to 64.9%.(Yeh *et al*. 1989; Liu *et al*. 2008)

A genome-wide association study (GWAS) examines characteristic genetic alterations and epigenetic profiles associated with various diseases and further detects the contribution of individual genetic factors to complex diseases, facilitating the identification of potential targets associated with the occurrence, development and treatment of a plethora of diseases.(Visscher *et al*. 2012) Here, we conducted a GWAS using human whole genome exon sequencing to analyze the genetic susceptibility of TP53 R249S and identify the genetic factors influencing the TP53 codon 249 mutation in AFB1 and HBV exposed patients with HCC in the Guangxi region.

## Methods

### Study participants

All patients with HCC subjected to hepatectomy from the Department of Hepatobiliary Surgery at the First Affiliated Hospital of Guangxi Medical University, China, from January 2001 to November 2013 were enrolled. For the replication stage study, 270 independent subjects were additionally recruited from December 2013 to August 2016. All cases were positive for hepatitis B surface antigen and HCC after hepatectomy, as confirmed via histopathology. The clinicopathological variables of enrolled participants, including age, gender, smoking status, drinking status, pathological grade, biobehavior of the cancer, serum AFP level, hepatic cirrhosis, radical resection and use of transcatheter hepatic arterial chemoembolization (TACE), were obtained from clinical records and pathological reports. Tumor status was classified according to the Barcelona Clinic Liver Cancer (BCLC) staging system.(Bruix and Sherman 2005) Child-Pugh classification was conducted as described previously.(Pugh *et al*. 1973) Portal vein tumor thrombus (PVTT) was determined according to previous criteria.(Kondo *et al*. 2009) Smoking status, drinking status and radical resection were defined as reported earlier.(Yu *et al*. 2016) All studies were approved by the Ethical Review Committee of the First Affiliated Hospital of Guangxi Medical University (Approval Number: 2015 [KY-E-072]). All participants signed an informed consent form.

### Specimens and TP53 codon 249 mutation detection

HCC samples were collected during surgery and immediately stored at −80°C. Total DNA was extracted using the TIANamp Genomic DNA Kit (TIANGEN BIOTECH [BEIJING] CO, LTD). DNA concentration and purity were measured with the NanoDrop2000 system (Thermo Fisher Scientific, Waltham, MA, USA). The TP53 R249S mutation was detected via Sanger sequencing after amplification with polymerase chain reaction (PCR). Sequencing primers are presented in **Table S1**.

### Immunohistochemistry

TP53 expression of HCC samples was evaluated using immunohistochemical staining as follows: antigen retrieval was conducted with EDTA-Tris at a high temperature for 2.5 min and slices treated with 3% hydrogen peroxide to inactivate endogenous peroxidase. After washing with PBS, slices were incubated with mouse antibody against human TP53 (1:150) (ZSGB-BIO ORIGENE, Beijing, China) at 4°C followed by secondary antibody (Dako Cytomation, Glostrup, Denmark) for 30 min at 37°C. Finally, slices were visualized with DAB (Dako Cytomation) and counterstained with hematoxylin. Positive and negative controls were included for each sample. All tissue sections were reviewed and scored blindly by two experienced pathologists. Where possible, a consensus of joint review was obtained in case of disagreement. The proportion of positive cells was calculated based on the following formula: (number of positive cells/total number of the cells×100%). Positive staining for TP53 was defined as ≥ 10% positive cancer cells.(Esrig *et al*. 1994; Cote *et al*. 1998)

### Genotyping and quality control in the GWAS

In the GWAS procedure, all samples were genotyped using Illumina HumanExome BeadChip-12-1_A, which includes 242,901 markers of protein-altering variants. The flow chart and genotyping procedure is depicted in **Figure S1** and **S2** respectively. Genotype calling was carried out using the Genotyping Module v1.0 in GenomeStudio version 2011.1, with an average call rate of 99.84%. A total of 50 samples (> 10%) were randomly selected and sequence analysis of candidate loci performed using ABI PRISM 3100 (Applied Biosystems, Shanghai Sangon Biological Engineering Technology & Services Co, Ltd, Shanghai, China). The results of sequencing were 100% concordant with those of genotyping using BeadChip-12-1A.

A quality control (QC) procedure was utilized before association analysis. 1. Samples were excluded under the following conditions: (i) genotyping rate < 95%, (ii) ambiguous gender, (iii) genome-wide identity-by-descent (IBD) > 0.1875, (iv) outliers in principal components analysis (PCA) for ancestry and population stratification. 2. SNPs were removed with a (i) genotype call rate < 95%, (ii) P value in Hardy-Weinberg equilibrium (HWE) of < 1 × 10^−6^, (iii) minor allele frequency (MAF) < 0.05. 3. PCA: analysis of population stratification by PCA was performed to eliminate multi-ethnic interference using the EIGENSOFT package. Genomic inflation factors (GIF) were used to investigate residual population stratification, which was calculated with MATLAB 7.0. The procedure was performed using Plink version 1.07, R 3.0.1 and EIGENSOFT package. After quality control, the total genotyping rate in the remaining individuals was determined, as shown in **Figure S3A**. Genotype failure rate and heterozygosity across all individuals are also presented in **Figure S3B**. The principal component analysis (PCA) plot disclosed no or mild population stratification in our study population (**Figure 1A**). The genomic control inflation factor was determined from a Quantile-Quantile (Q-Q) plot (**Figure 1B**). Ultimately, 21,501 SNPs from 459 cases were used for the GWAS after quality control.

**Figure 1.**
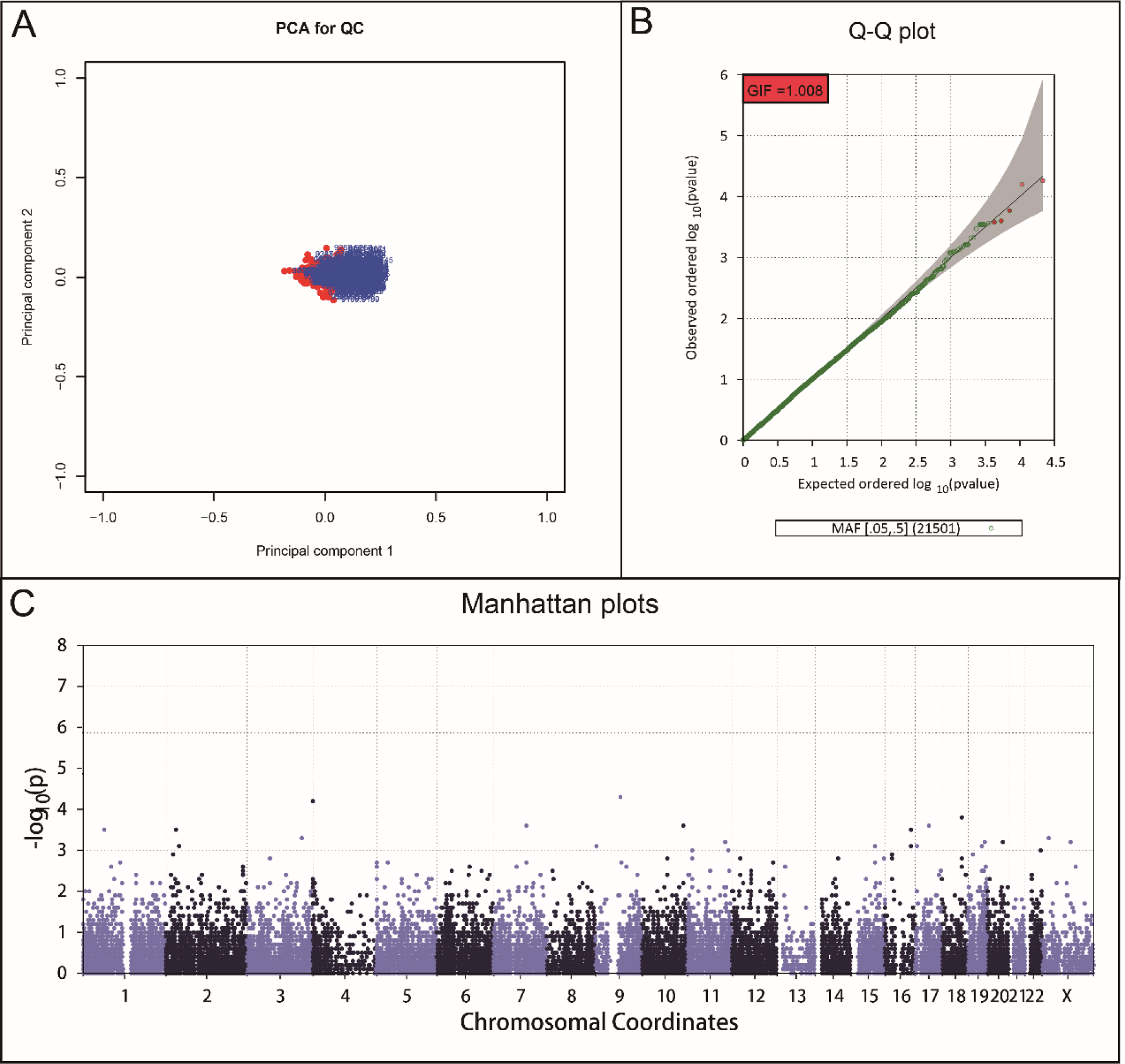
The results of genome-wide association analysis. A, The results of principal component analysis across all individuals. B, Quantile-Quantile (Q-Q) plots from *P* value of Single Variant Test. C, Manhattan plots for association analysis. Results from Single Variant Test (−log10 P values) were plotted against genomic position (GRCh37/hg19).

### SNP selection for the replication study

Based on the presence or absence of TP53 R249S, subjects were classified into mutation and non-mutation groups. Genetic differences between the two groups were explored using the GWAS. The EPACTS package was used to perform the association analysis. According to Manhattan plot (**Figure 1C**), 35 SNPs with top-ranking P values were selected. However, only 29 primer pairs could be designed for these SNPs to perform the replication study. The 29 SNPs used for validation analyses are specified in **Table S2** and their corresponding primers in **Table S3**.

### Genotyping and quality control in the replication study

In replication stage, total 270 subjects were enrolled. Genotyping analyses used iPLEX Gold SNP typing technology based on the MassARRAY® array platform. The quality control procedure was as follows: 1. samples were excluded based on (i) genotyping rate of < 95% and (ii) genome-wide IBD > 0.1875, 2. SNPs were removed that showed (i) a genotype call rate <95%, (ii) P value in HWE < 1 × 10^−6^ and (iii) MAF < 0.05. Following quality control, 258 samples (80 in the mutation and 178 in the non-mutation group) were included for analysis. The results of the replication study on 29 SNPs are detailed in **Table S4** and the corresponding primers presented in **Table S5**.

### Quantitative RT-PCR

Tumor and paired non-tumor liver tissue samples were collected for gene expression analysis from 20 subjects with HCC randomly selected from the independent replication study. Candidate SNPs were selected from the GWAS and replication analyses to analyze potential associations with their located gene expression. RNA was extracted from samples using TRIzol and cDNA synthesized from 1 μg RNA template using PrimeScript™ RT reagent Kit with gDNA Eraser (TAKARA BIO INC). The sequences of primers used for amplification are shown in Table S6. Quantitative RT-PCR was conducted in triplicate on an ABI7500 Real-Time PCR System using FastStart Universal SYBR Green Master (ROX). Expression of all samples was normalized to that of GAPDH using the ΔC_T_ method.

### Statistical analysis

Results are expressed as means ± SD for normally distributed variables or median for non-normal distribution. The logistic regression model was applied to analyze differences in clinicopathological factors between R249S mutation and non-mutation groups. OR and 95% CI values were calculated in univariate analysis. Analysis of associations between TP53 R249S and SNPs was performed using Single Variant Test 68 (Logistic Score Test) with EPACTS package version 3.2.6(Lin and Tang 2011) and quantile-quantile and Manhattan plots generated. The Chi-squared test and logistic regression model were applied to determine the association of genetic models with gene types. Local linkage disequilibrium (LD) and recombination patterns were analyzed using LocusZoom (see URL).(Pruim *et al*. 2010) Differences in expression between tumors and paired adjacent non-tumor samples were assessed with Student’s t-test, and differences between SNP genotypes and gene expression with a non-parametric trend test. Statistical analysis was performed with SPSS version 18.0 (SPSS, Chicago, IL, USA). Data were considered significant at P < 0.05.

## Results

### Baseline Characteristics

In the GWAS, 165 mutation and 320 non-mutation cases were analyzed while the replication study included 80 mutation and 178 non-mutation samples. No statistical differences in clinicopathological parameters were evident between the subgroups stratified by TP53 R249S mutation, including age, gender, race, BMI, smoking status, drinking status, Child-Pugh score, BCLC stage, TACE status, cirrhosis, serum alpha fetoprotein (AFP) level, pathological grade, antiviral therapy, tumor size, capsule, regional invasion, intrahepatic metastasis, vascular invasion and PVTT in both stages, except for number of tumors in the replication study (*P* < 0.049, HR = 2.72, 95% CI = 1.00–7.38; **Table 1 and Table 2**). Notably, however, TP53 expression status was significantly different between the mutation and non-mutation groups in GWAS (*P* < 0.001, OR = 4.46, 95% CI = 2.73–7.27; **Table 3**) and in replication stage analysis (*P* = 0.006, OR = 3.15, 95% CI = 1.40–7.09; **Table 3**).

**Table 1.**
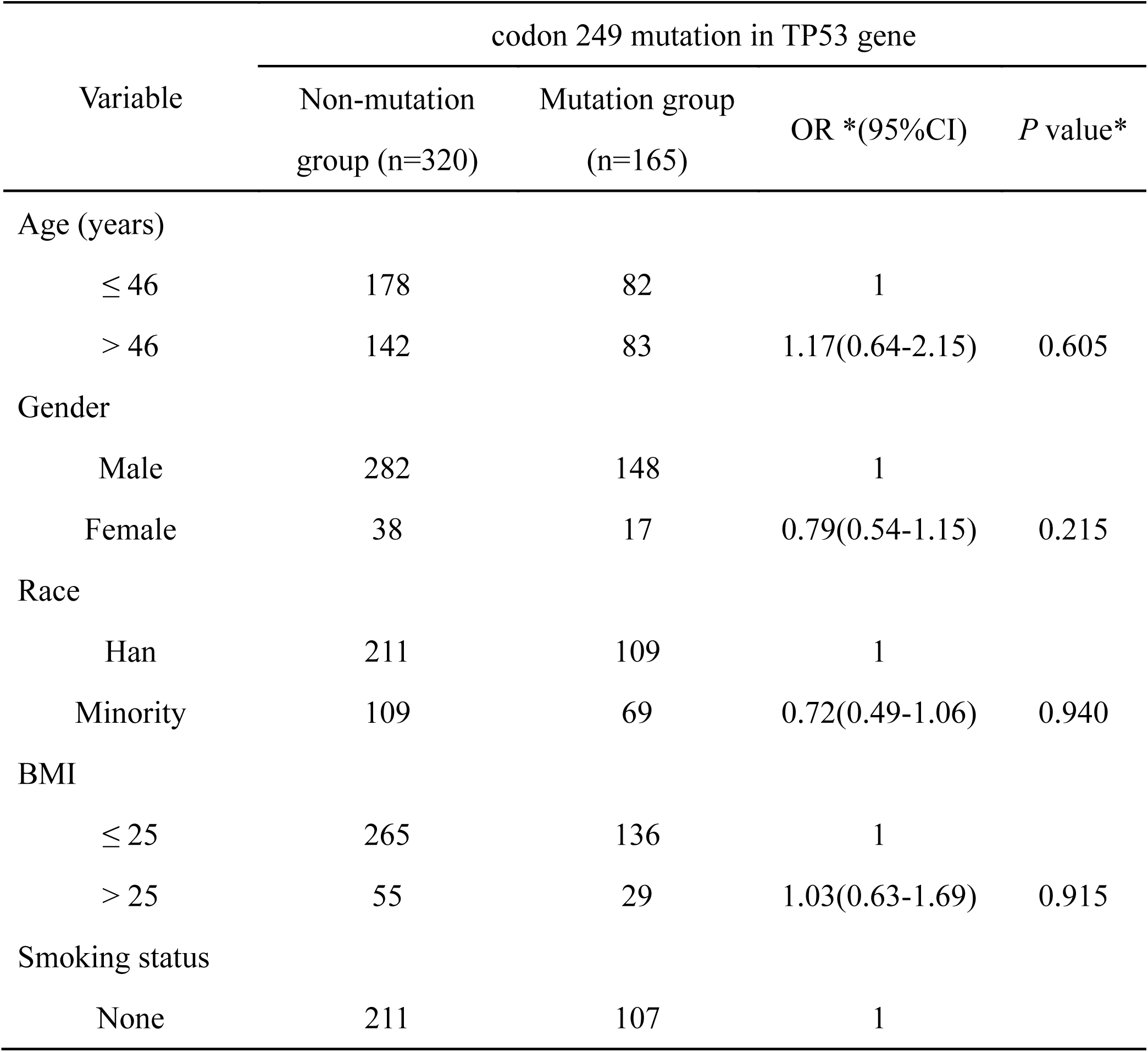
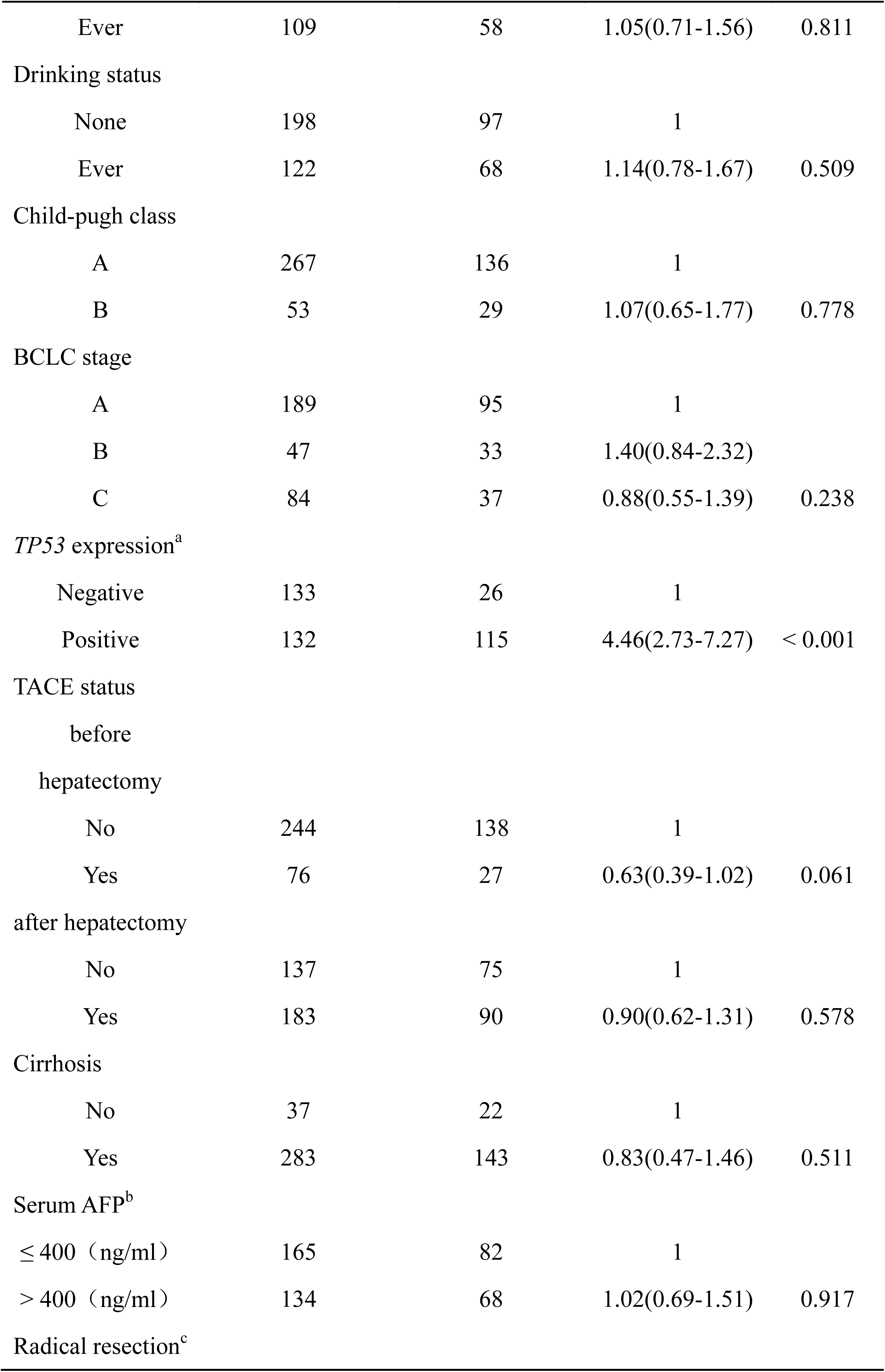
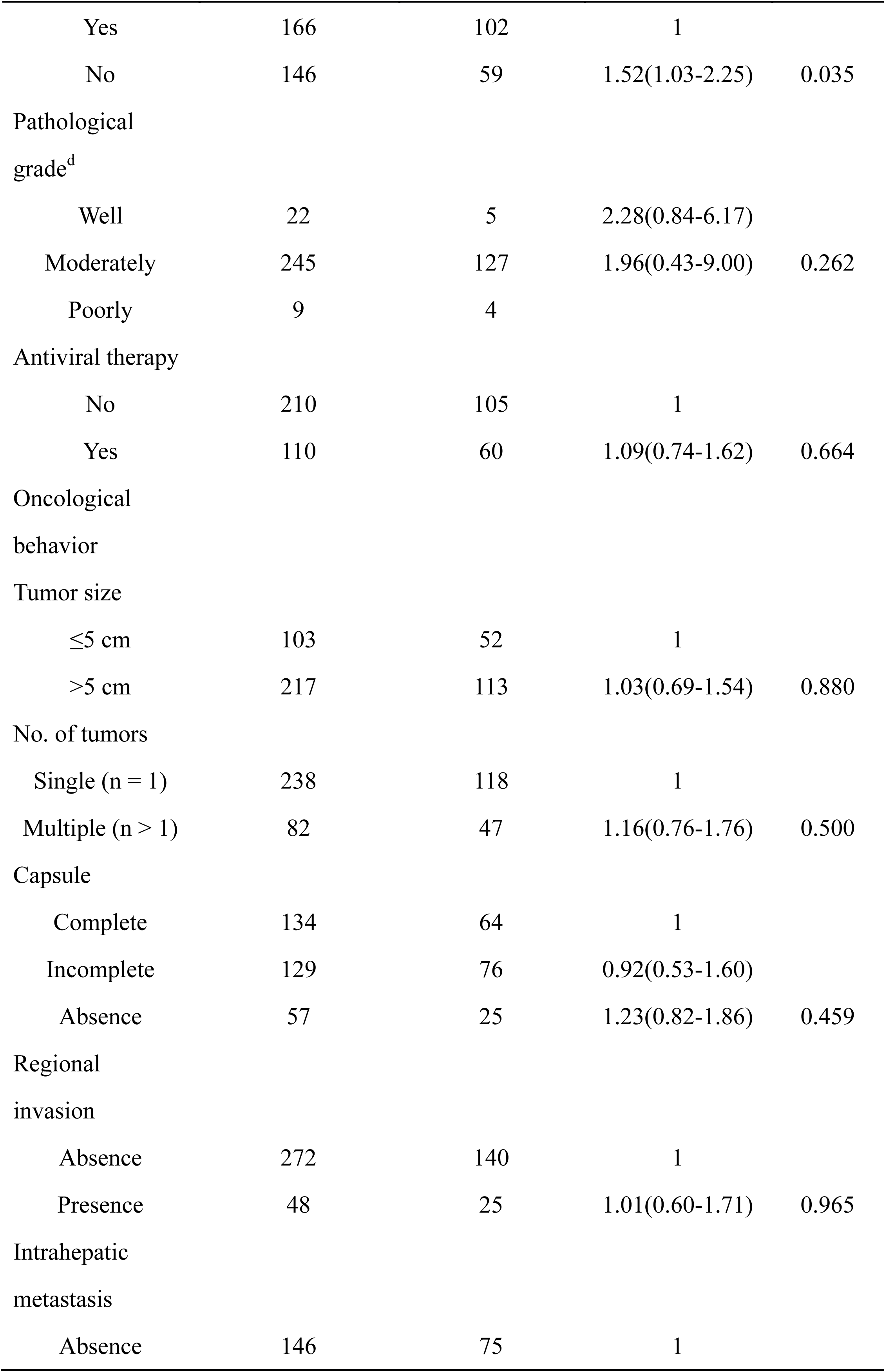
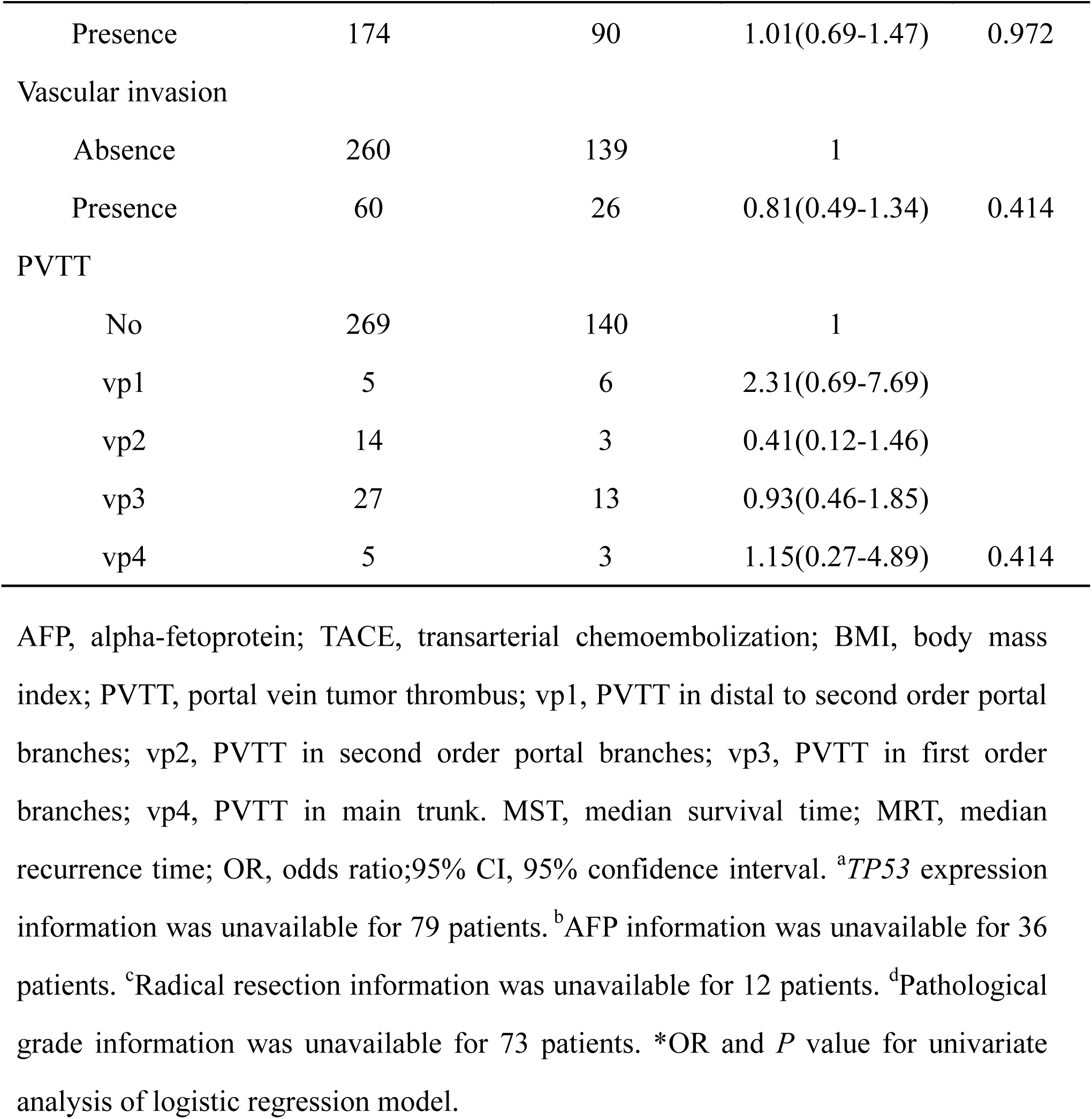
Clinicopathological characteristics of subjects analyzed in the GWAS stage

**Table 2.**
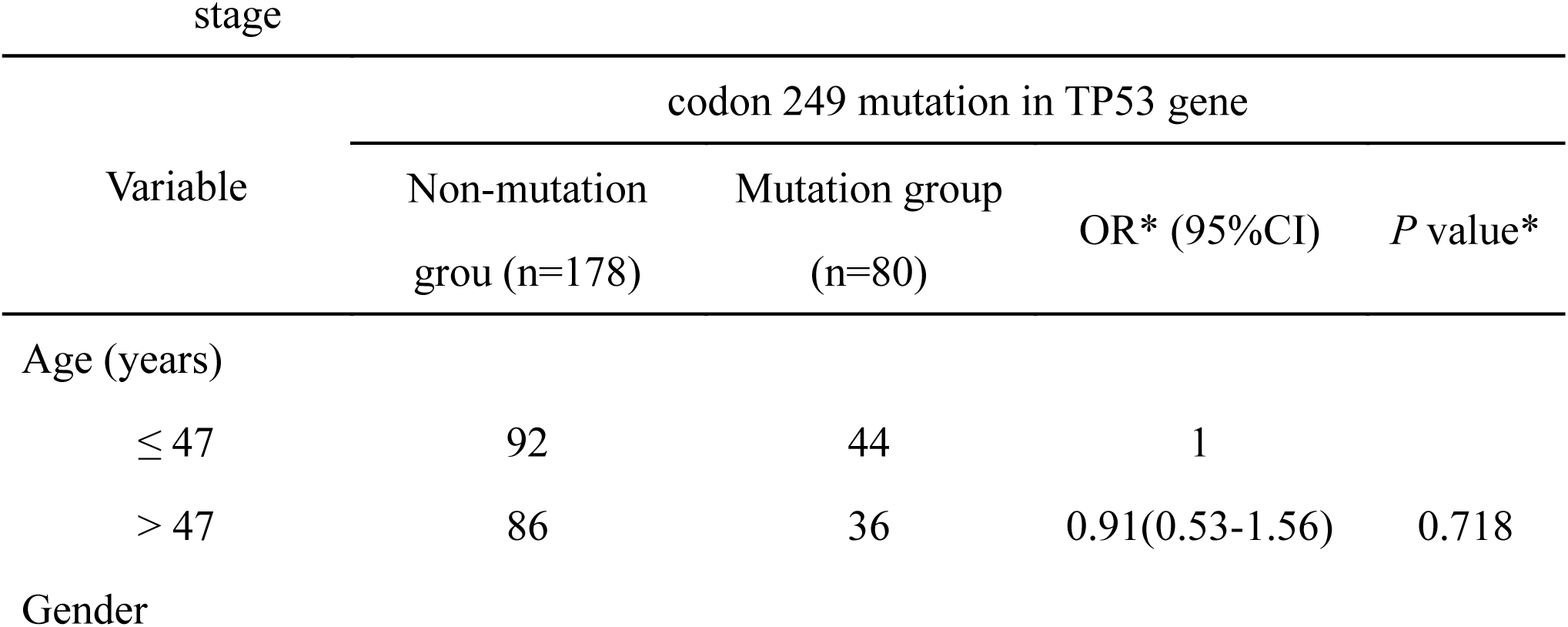
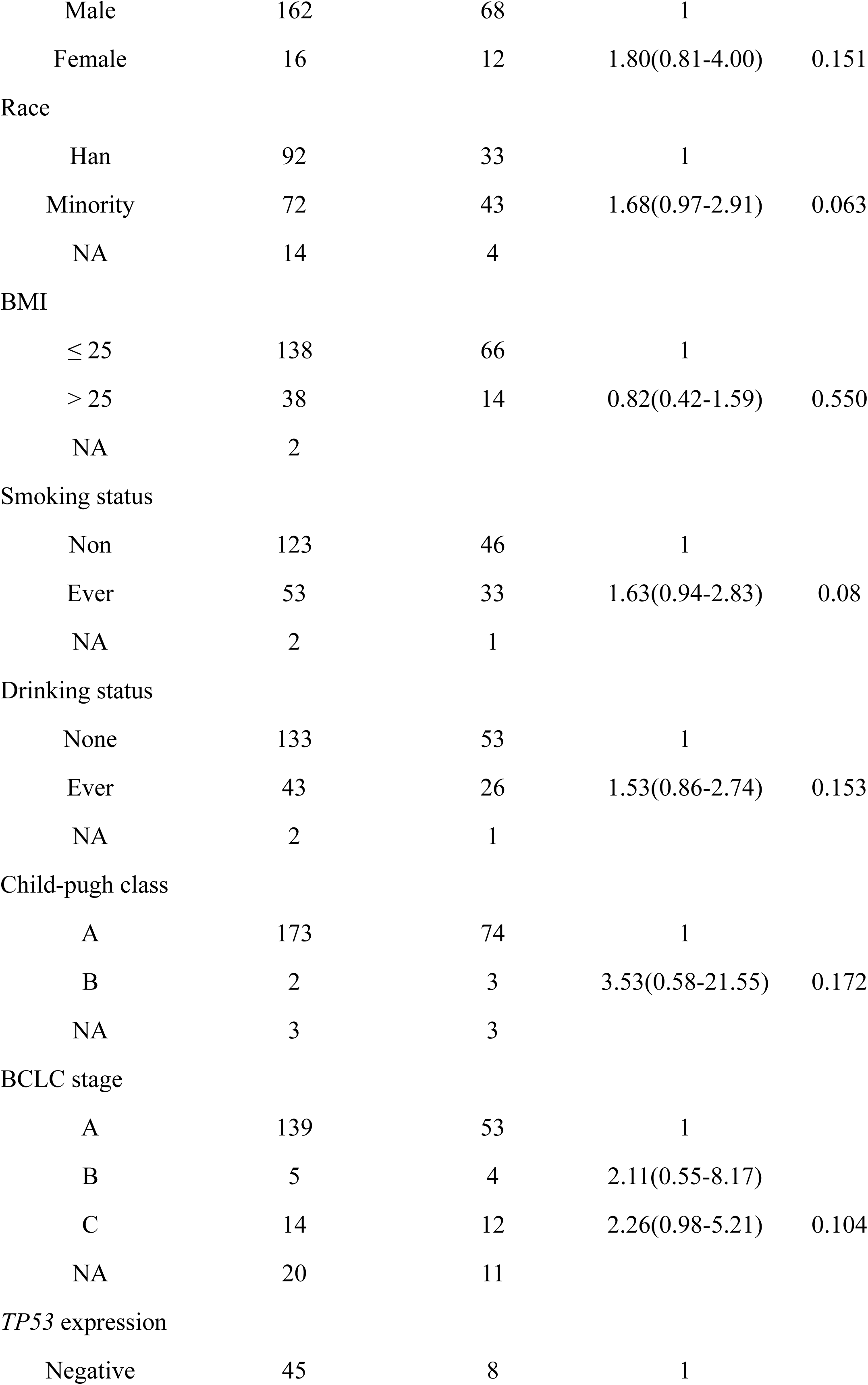
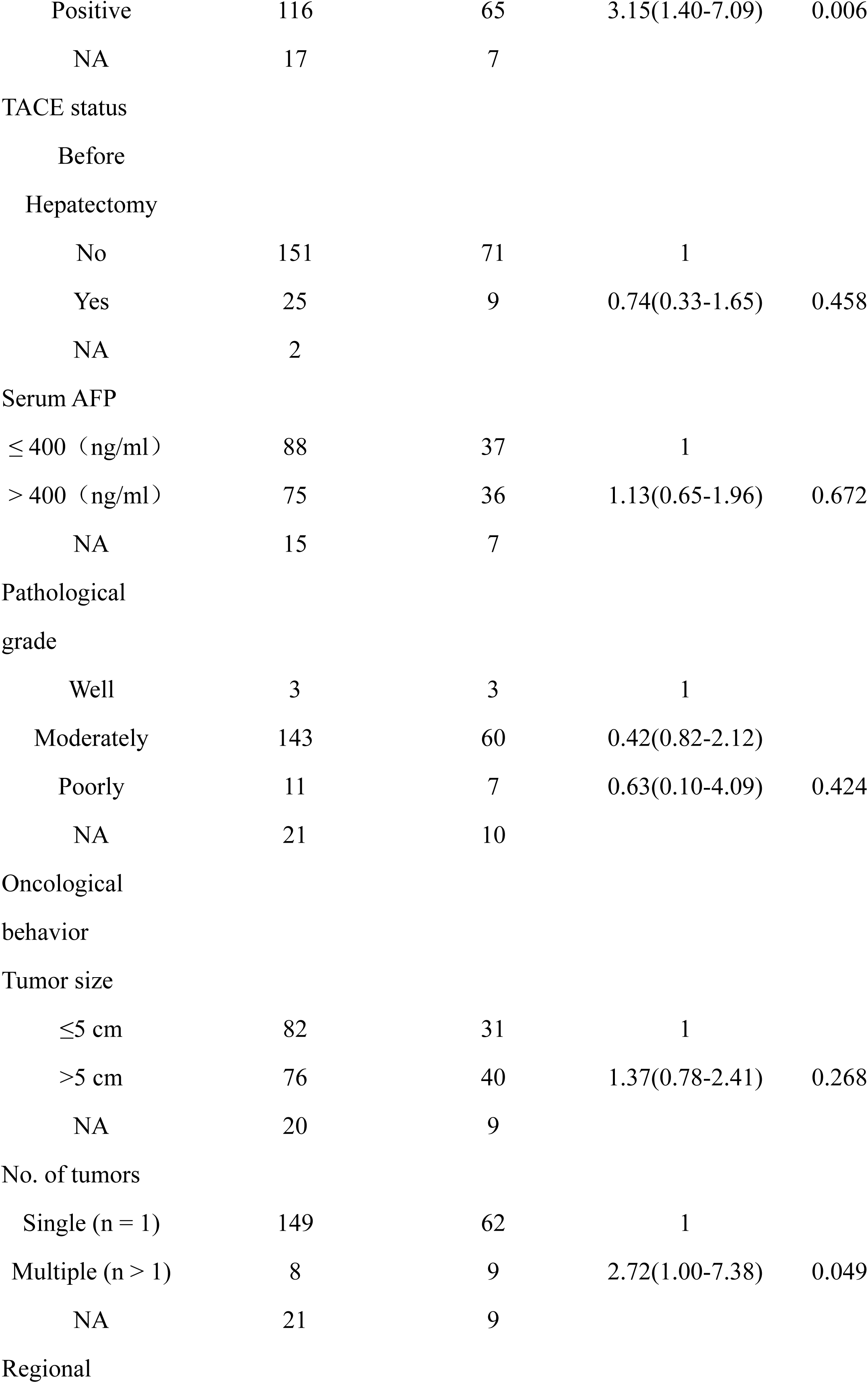
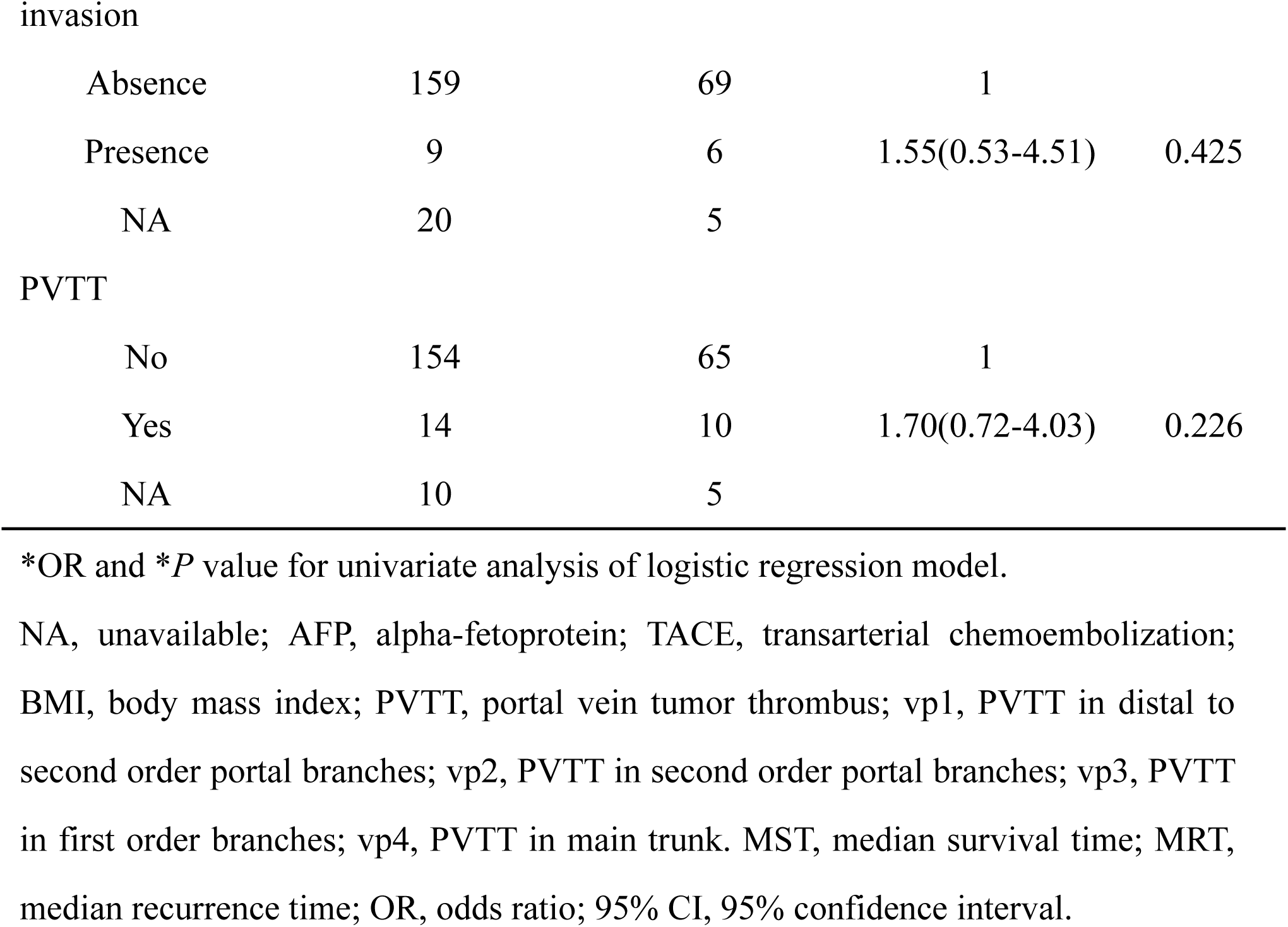
Clinicopathological characteristics of subjects analyzed in the replication invasion

| Variable | codon 249 mutation in TP53 gene |  |  |  |
| --- | --- | --- | --- | --- |
|  | Non-mutation<br>group (n=178) | Mutation group<br>(n=80) | OR* (95%CI) | *P* value* |
| Age (years) |  |  |  |  |
| ≤ 47 | 92 | 44 | 1 |  |
| > 47 | 86 | 36 | 0.91(0.53-1.56) | 0.718 |
| Gender |  |  |  |  |
| Male | 162 | 68 | 1 |  |
| Female | 16 | 12 | 1.80(0.81-4.00) | 0.151 |
| Race |  |  |  |  |
| Han | 92 | 33 | 1 |  |
| Minority | 72 | 43 | 1.68(0.97-2.91) | 0.063 |
| NA | 14 | 4 |  |  |
| BMI |  |  |  |  |
| ≤ 25 | 138 | 66 | 1 |  |
| > 25 | 38 | 14 | 0.82(0.42-1.59) | 0.550 |
| NA | 2 |  |  |  |
| Smoking status |  |  |  |  |
| Non | 123 | 46 | 1 |  |
| Ever | 53 | 33 | 1.63(0.94-2.83) | 0.08 |
| NA | 2 | 1 |  |  |
| Drinking status |  |  |  |  |
| None | 133 | 53 | 1 |  |
| Ever | 43 | 26 | 1.53(0.86-2.74) | 0.153 |
| NA | 2 | 1 |  |  |
| Child-pugh class |  |  |  |  |
| A | 173 | 74 | 1 |  |
| B | 2 | 3 | 3.53(0.58-21.55) | 0.172 |
| NA | 3 | 3 |  |  |
| BCLC stage |  |  |  |  |
| A | 139 | 53 | 1 |  |
| B | 5 | 4 | 2.11(0.55-8.17) |  |
| C | 14 | 12 | 2.26(0.98-5.21) | 0.104 |
| NA | 20 | 11 |  |  |
| TP53 expression |  |  |  |  |
| Negative | 45 | 8 | 1 |  |
| Positive | 116 | 65 | 3.15(1.40-7.09) | 0.006 |
| NA | 17 | 7 |  |  |
| TACE status |  |  |  |  |
| Before |  |  |  |  |
| Hepatectomy |  |  |  |  |
| No | 151 | 71 | 1 |  |
| Yes | 25 | 9 | 0.74(0.33-1.65) | 0.458 |
| NA | 2 |  |  |  |
| Serum AFP |  |  |  |  |
| ≤ 400 (ng/ml) | 88 | 37 | 1 |  |
| > 400 (ng/ml) | 75 | 36 | 1.13(0.65-1.96) | 0.672 |
| NA | 15 | 7 |  |  |
| Pathological |  |  |  |  |
| grade |  |  |  |  |
| Well | 3 | 3 | 1 |  |
| Moderately | 143 | 60 | 0.42(0.82-2.12) |  |
| Poorly | 11 | 7 | 0.63(0.10-4.09) | 0.424 |
| NA | 21 | 10 |  |  |
| Oncological |  |  |  |  |
| behavior |  |  |  |  |
| Tumor size |  |  |  |  |
| ≤5 cm | 82 | 31 | 1 |  |
| >5 cm | 76 | 40 | 1.37(0.78-2.41) | 0.268 |
| NA | 20 | 9 |  |  |
| No. of tumors |  |  |  |  |
| Single (n = 1) | 149 | 62 | 1 |  |
| Multiple (n > 1) | 8 | 9 | 2.72(1.00-7.38) | 0.049 |
| NA | 21 | 9 |  |  |
| Regional |  |  |  |  |
| Absence | 159 | 69 | 1 |  |
| Presence | 9 | 6 | 1.55(0.53-4.51) | 0.425 |
| NA | 20 | 5 |  |  |
| PVTT |  |  |  |  |
| No | 154 | 65 | 1 |  |
| Yes | 14 | 10 | 1.70(0.72-4.03) | 0.226 |
| NA | 10 | 5 |  |  |

\*OR and \*P value for univariate analysis of logistic regression model.
NA, unavailable; AFP, alpha-fetoprotein; TACE, transarterial chemoembolization; BMI, body mass index; PVTT, portal vein tumor thrombus; vp1, PVTT in distal to second order portal branches; vp2, PVTT in second order portal branches; vp3, PVTT in first order branches; vp4, PVTT in main trunk. MST, median survival time; MRT, median recurrence time; OR, odds ratio; 95% CI, 95% confidence interval.

**Table 3.** Analysis of the R249S mutation in relation to TP53 expression in the GWAS and replication stage experiments.

| Subjects | Number | TP53 R249S mutation |  |  |  |
| --- | --- | --- | --- | --- | --- |
|  |  | Non-mutation group(n) | Mutation group (n) | OR* (95%CI) | *P* value* |
| GWAS |  |  |  |  |  |
| Negative | 159 | 133 | 26 | 1 |  |
| Positive | 247 | 132 | 115 | 4.46(2.73–7.27) | **< 0.001** |
| NA | 79 | 55 | 24 |  |  |
| Replication |  |  |  |  |  |
| Negative | 53 | 45 | 8 | 1 |  |
| Positive | 181 | 116 | 65 | 3.15(1.40–7.09) | **0.006** |
| NA | 24 | 17 | 7 |  |  |

OR, odds ratio; 95% CI, 95% confidence interval
\*OR and P value for univariate analysis of the logistic regression model

### Quality control (QC)

After QC, a total of 459 samples with 21,501 SNPs were identified and the total genotyping rate in the remaining individuals was 98.35% (**Figure S3A**). Data from the PCA plot demonstrated no or mild stratification in our study population (**Fig 1A**). The Q-Q plot revealed a genomic control inflation factor (λ) of 1.008 (**Figure 1B**).

### Association Analysis

Overall, 258 samples met the experimental requirements for the replication study. As shown in **Table 4**, the SNPs rs8022091, rs9930984, rs75218075 were significantly associated with TP53 R249S at the GWAS stage (MAF = 0.36, *P* = 0.00155; MAF = 0.126, *P* = 0.000334; MAF = 0.179, *P* = 0.00047), replication stage (MAF = 0.347, *P* = 0.0232; MAF = 0.091, *P* = 0.0255; MAF = 0.147, *P* = 0.0295) and combined analyses (MAF = 0.356, *P* = 0.042; MAF = 0.113, *P* = 4.84 × 10^−6^; MAF = 0.167, *P* = 7.36 × 10^−5^) after adjustment for age, sex, race, smoking and drinking status. Ultimately, rs8022091, rs9930984, rs75218075 were identified as SNPs associated with TP53 R249S mutation and subjected to further analyses.

**Table 4.**
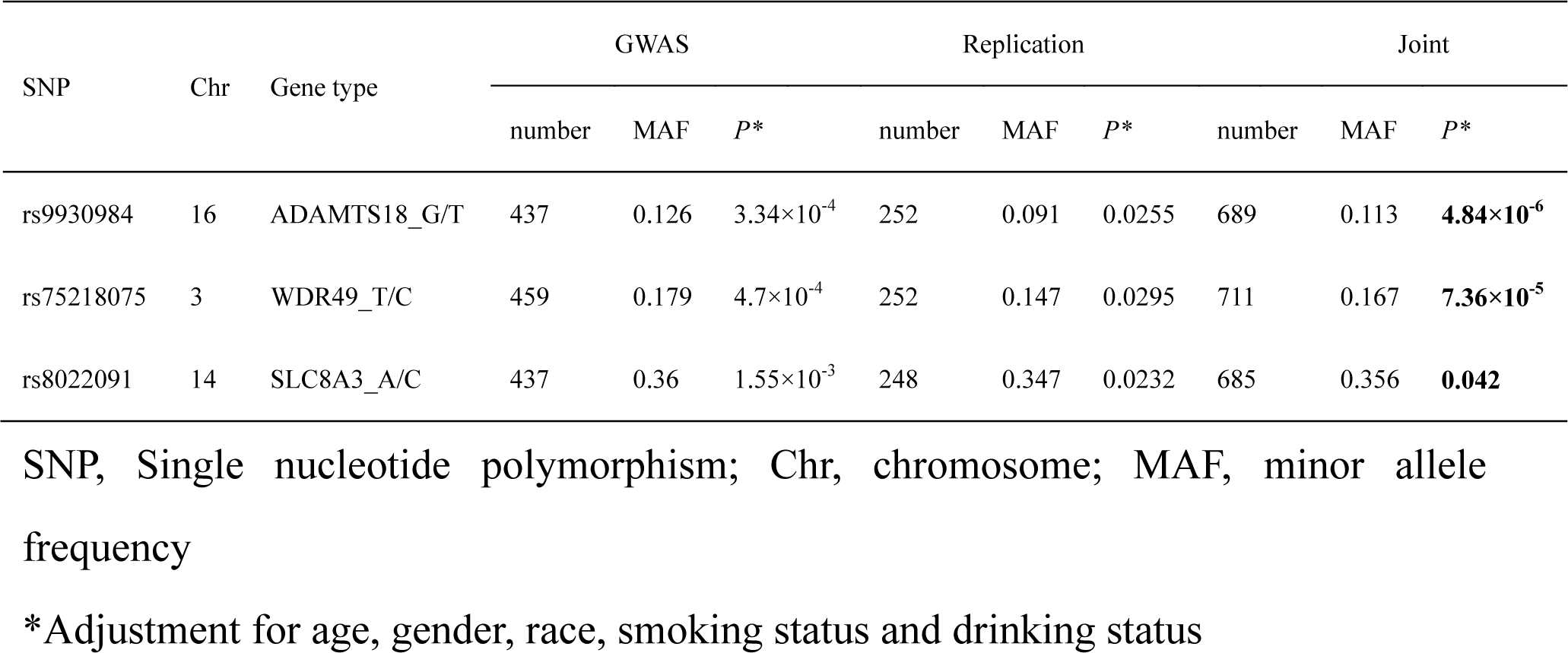
Association analysis of candidate SNPs in GWAS, replication stage and combined studies.

### Genetic model analysis

To establish the potential association of genotype variants of candidate SNPs with TP53 R249S mutation, additive and dominant genetic models were constructed and results listed in **Table 5**. In the additive genetic model, TP53 R249S was significantly associated with the TT genotype at rs9930984 (*P* = 0.010, OR = 4.94, 95% CI = 1.48– 16.52) and rs75218075 (*P* = 0.002, OR = 5.79, 95% CI = 1.33–25.13). In the dominant genetic model, populations with HCC carrying the TT genotype of rs9930984 showed greater risk of TP53 R249S, compared with those carrying GG and GT genotypes (*P* = 6.43 × 10^−5^, OR = 2.27, 95% CI = 1.67–4.44). A similar result was obtained with the population carrying TT genotype of rs75218075, which was associated with higher risk of TP53 R249S, relative to populations carrying CC and CT genotypes (*P* = 2.16 × 10^−4^, OR = 1.98, 95% CI = 1.38–2.85). In contrast, the rs8022091 genotypes did not demonstrate statistical significance between the mutation and non-mutation groups in both the additive and dominant genetic models.

**Table 5.** Combined analysis for genotypes of associated SNPs and the *TP53* R249S mutation

| SNPs | TP53 R249S mutation |  | OR (95% CI) | *P* | Test power |
| --- | --- | --- | --- | --- | --- |
|  | Non-mutation | Mutation |  |  |  |
| rs9930984 |  |  |  |  |  |
| Additive |  |  |  |  | 0.85 |
| GG | 26 | 3 | 1 | $3.06 \times 10^{-4}$ | |
| GT | 79 | 19 | 2.08 (0.57–7.62) | 0.267 |  |
| TT | 358 | 204 | 4.94 (1.48–16.52) | **0.010** |  |
| Dominant |  |  |  |  | 0.84 |
| GG+GT | 105 | 22 | 1 |  |  |
| TT | 358 | 204 | 2.27(1.67–4.44) | $6.43 \times 10^{-5}$ | |
| rs75218075 |  |  |  |  |  |
| Additive |  |  |  |  | NA |
| CC | 19 | 2 | 1 | **0.001** |  |
| CT | 147 | 49 | 3.17 (0.71–14.89) | 0.130 |  |
| TT | 307 | 187 | 5.79(1.33–25.13) | **0.002** |  |
| Dominant |  |  |  |  | 0.65 |
| CC+CT | 166 | 51 | 1 |  |  |
| TT | 307 | 187 | 1.98(1.38–2.85) | **2.16×10<sup>-4</sup>** |  |
| rs8022091 |  |  |  |  |  |
| Additive |  |  |  |  | 0.82 |
| AA | 65 | 40 | 1 | 0.413 |  |
| AC | 185 | 92 | 0.81(0.51–1.29) | 0.371 |  |
| CC | 209 | 94 | 0.73(0.46–1.16) | 0.184 |  |
| Dominant |  |  |  |  | 0.79 |
| AA | 65 | 40 | 1 |  |  |
| AC+CC | 394 | 186 | 0.77(0.50-1.18) | 0.228 |  |

SNP = single nucleotide polymorphism; OR, odds ratio; CI = confidence interval *P* values, OR and 95% CI were calculated under the additive model via logistic regression while adjusting for age, sex, race, smoking status and drinking status.

### LD and Haplotype Analysis

Analysis was performed for LD and haplotypes about 2 Mb nearby, specifically, rs9930984 located in ADAM metallopeptidase with thrombospondin type 1 motif 18 (ADAMTS18) on chromosome 16, rs75218075 located in the WD repeat domain 49 (WDR49) on chromosome 3 and rs8022091 located in solute carrier family 8 member A3 (SLC8A3) on chromosome 14 (**Figure** 2A–2C). Our data suggested that rs9930984 is in LD with rs11640912. However, no significant association was evident between rs11640912 and TP53 R249S in the replication stage analysis (**Table S4**). And rs8022091 and rs75218075 did not find any linkage disequilibrium SNP locus nearby.

### Quantitative RT-PCR

To establish the effects of genotypes at the three SNP sites on their located gene, mRNA expression of ADAMTS18, WDR49 and SLC8A5 was assessed. ADAMTS18 expression was significantly higher in HCC tissue, compared with that in paired non-tumor tissue (*P* =0.041, **Figure 3A**) and populations with the TT genotype at rs9930984 showed lower expression in non-tumor tissue relative to individuals with the GT genotype (*P* = 0.0028, **Figure 3A**). However, no significant differences in ADAMTS18 expression were detected between GT and TT genotypes of rs9930984 in HCC, TP53 R249S mutation and non-mutation groups of HCC or non-tumor tissue (**Figure 3A**). WDR49 expression was markedly lower in HCC relative to paired non-tumor tissue (*P* = 0.0011). Similarly, we observed no significant differences in WDR49 expression between the genotypes of rs75218075, TP53 R249S mutation and non-mutation groups of HCC or non-tumor tissue (**Figure 3B**). SLC8A5 expression was not significantly detected in HCC and paired non-tumor tissue, diverse genotypes of rs75218075, TP53 R249S mutation and non-mutation groups or non-tumor tissue (**Figure 3C**).

**Figure 2.**
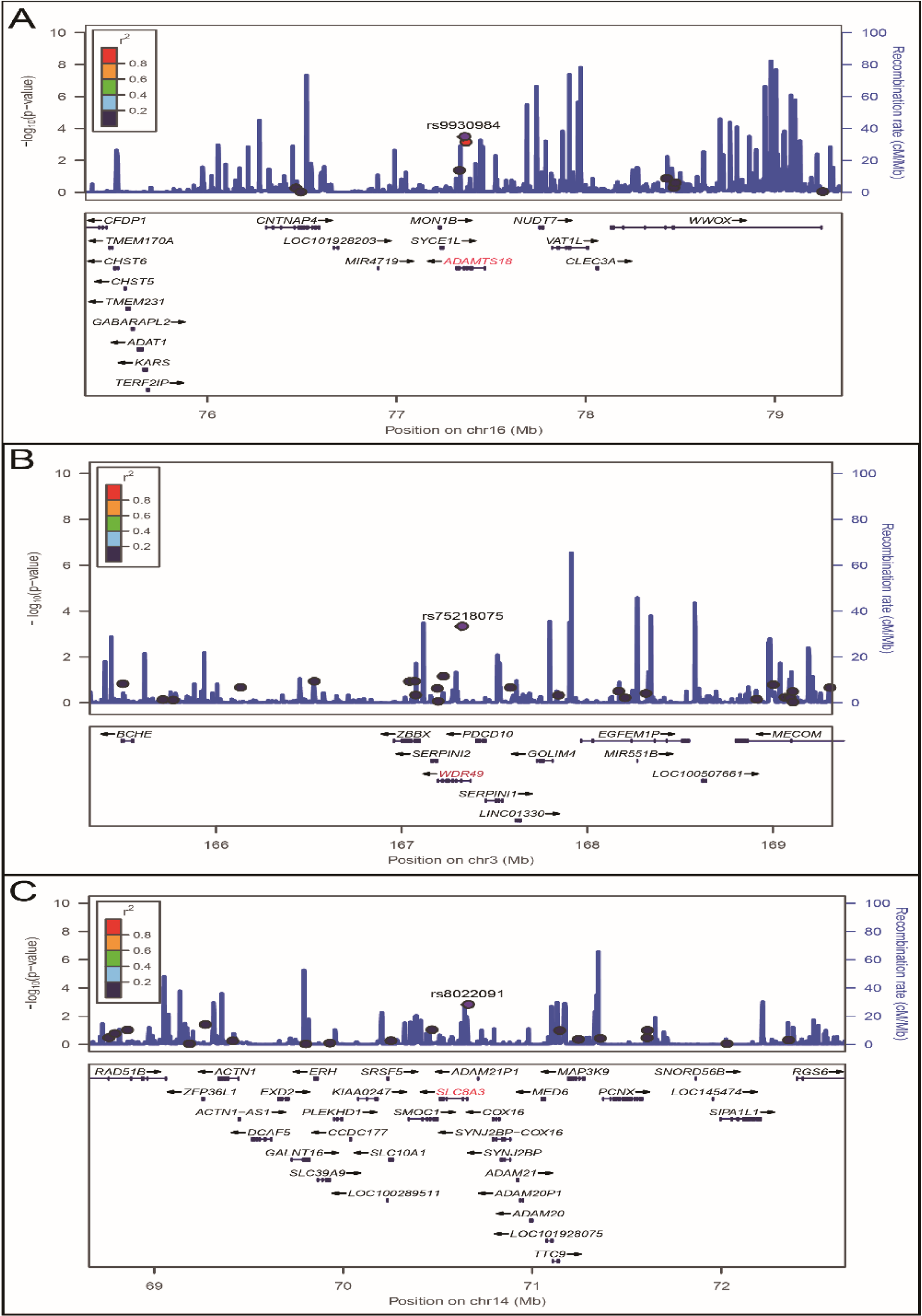
Locus Zoom plot analysis of local linkage disequilibrium (LD) for candidate SNPs. A, Locus Zoom plot for rs8022091 B, Locus Zoom plot for rs9930984 C, Locus Zoom plot for rs752180.

**Figure 3.**
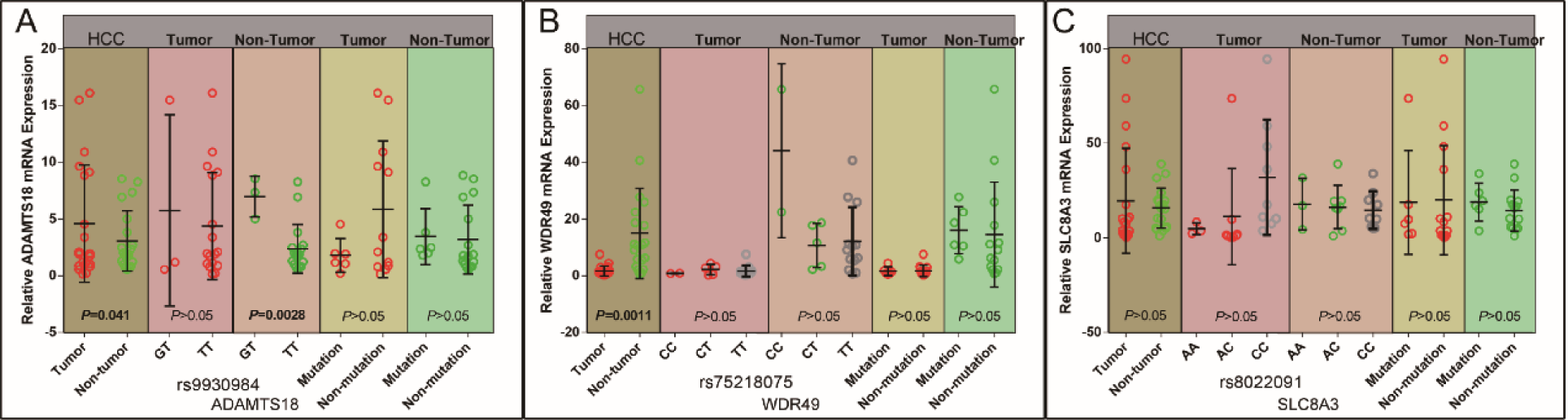
Relative expression of candidate genes in HCC. A, Relative expression of ADAMTS18 in HCC and non-tumor tissue, tumor or non-tumor tissue with and without the TP53 R249S mutation or according to rs9930984 genotype. B, Relative expression of WDR49 in HCC and non-tumor tissue, tumor or non-tumor tissue with or without the TP53 R249S mutation or according to rs75218075 genotype. C, Relative expression of SLC8A3 in HCC and non-tumor tissue, tumor or non-tumor tissue with or without the TP53 R249S mutation or according to rs8022091 genotype.

### Discussion

To identify novel susceptibility loci for the TP53 R249S mutation in HCC with AFB1 and HBV exposure, we performed a genome-wide association study (GWAS). A trend of positive TP53 expression was evident in HCC with the R249S mutation, compared with HCC without R249S in both GWAS and replication stage analysis which was consistent with previous findings.(Qi *et al*. 2015) Following a combined analysis of all subjects from the GWAS and the subsequent replication study, rs8022091, rs9930984 and rs75218075 remained significantly associated with TP53 R249S after adjustment for age, gender, race, smoking status and drinking status.

We further conducted a combined analysis for genotypes of these three SNPs and TP53 R249S. Our results suggest that the TT genotype of rs9930984 and rs75218075 are risk factors associated with incidence of the TP53 R249S mutation in HCC coupled with AFB1 and HBV exposure. Consistently, TT genotypes of rs9930984 and rs75218075 were significantly associated with TP53 R249S in HCC in both the additive genetic model. In contrast, genotypes of rs8022091 did not demonstrate significant associations with TP53 R249S in either the additive or dominant model. We performed further analysis for LD across 2 Mb nearby rs8022091, rs9930984 and rs75218075, which indicated that rs9930984 is in strong LD with rs11640912. However, rs11640912 was not significantly associated with TP53 R249S in our replication stage analysis.

rs9930984 is located in the ADAMTS18 gene, which encodes a member of the ADAMTS (disintegrin and metalloproteinase with thrombospondin motif) protein family. ADAMTS18 has been identified as a novel functional tumor suppressor.(Jin *et al*. 2007) Accumulating studies(Jin *et al*. 2007; Nordgard *et al*. 2008; Li *et al*. 2010; Wei *et al*. 2010; Wagstaff *et al*. 2011) have confirmed that mutations in this gene and hypermethylation of promoters are closely related to growth of a variety of tumors, suggesting a role in tumor suppression. Compared with paired adjacent or normal tissues, ADAMTS18 CpG methylation is significantly upregulated in gastric, colorectal and pancreatic cancers.(Li *et al*. 2010) Pharmacological and genetic demethylation analyses indicate that CpG methylation directly inhibits expression of ADAMTS18. In vitro, restoration of ADAMTS18 gene expression downregulated or silenced in tumor cell lines inhibits tumor cell clonality.(Jin *et al*. 2007) However, in a study on malignant melanoma, mutated ADAMTS18 was shown to promote cell growth, migration and metastasis.(Wei *et al*. 2010) In our study, ADAMTS18 rs9930984 was significantly correlated with the TP53 R249S mutation and populations carrying the TT genotype of rs9930984 showed higher risk of TP53 R249S, compared with those carrying the GG and GT genotypes. Results from separate experiments disclosed significantly higher ADAMTS18 mRNA expression in HCC relative to non-tumor tissue. However, no differences in ADAMTS18 expression were observed between the TP53 R249S mutation and non-mutation groups in both tumor and non-tumor tissue samples. Expression of ADAMTS18 in non-tumor samples carrying the TT genotype of rs9930984 was lower than that in samples with the GT genotype. To our knowledge, no studies to date have demonstrated associations of ADAMTS18 with HCC pathogenesis or the TP53 R249S mutation. Moreover, no relevant research on rs9930984 and its potential function is documented in the literature.

rs75218075 is located in the WDR49 gene on chromosome 3, which encodes a member of the protein family with nine WD repeats. These proteins are involved in multiple cellular processes, including cell cycle progression, signal transduction, apoptosis and gene regulation.(Neer *et al*. 1994; Smith *et al*. 1999) WDR79 is reported to be involved in the development and progression of non-small cell lung cancer.(Sun *et al*. 2016) An earlier study on bladder cancer by Chen et al.(Chen *et al*. 2015) revealed that WDR5 is upregulated in tumor tissues, in turn, promoting proliferation of bladder cancer cells, autologous renewal of cells and increased chemoresistance in vitro. However, no reports on the biological function of WDR49 have been presented until now and its role in tumor development remains to be established. Data from the present study indicate that WDR49 rs75218075 is significantly associated with the TP53 R249S mutation (Table 4, Tables S2 and S4) and patients carrying the TT genotype of rs75218075 are potentially at greater risk of TP53 R249S, compared with those carrying the CC and CT genotypes. WDR49 mRNA expression was significantly lower in HCC, compared with non-tumor tissue, but not correlated with genotypes of rs75218075, TP53 R249S mutation and non-mutation groups in HCC or non-tumor tissue.

rs8022091 is located in the SLC8A gene on chromosome 14 encoding a member of the sodium/calcium exchanger integral membrane protein family. Recent studies have shown that SLC8A3 is essential for activation of PKCα by Ca^2+^ in endothelial cell membrane, a necessary step for VEGF-induced ERK1/2 phosphorylation and angiogenesis.(Andrikopoulos *et al*. 2011a; Andrikopoulos *et al*. 2011b) Experiments on other cell types have demonstrated reverse transport of SLC8A3 in cardiac fibroblasts (Andrikopoulos *et al*. 2011a) and neuroblastoma cells,(Nashida *et al*. 2011) which is required for ERK1/2 activation. Another recent study showed a significant negative correlation between promoter CpG island methylation and SLC8A3 gene expression in samples of clear-cell renal cell cancer,(Deckers *et al*. 2017) indicating potential functional epigenetic control for the SLC8A3 gene in tumors. In both GWAS and replication experiments in the current study, SLC8A3 rs8022091 was significantly associated with the TP53 R249S mutation, while no correlation of genotypes of rs8022091 with the codon 249 mutation was observed. In a separate validation study, no significant differences in SLC8A3 mRNA expression were detected between HCC and non-tumor tissues, along with no evidence of association with genotypes of rs8022091, TP53 R249s mutation and non-mutation groups in HCC or non-tumor tissue.

Our comprehensive GWAS and replication stage studies on HCC with AFB1 and HBV exposure in the Chinese population were successfully conducted. ADAMTS18 rs9930984 and WDR49 rs75218075 were clearly associated with susceptibility to TP53 R249S and TT genotypes of rs9930984 and rs75218075 further identified as risk factors for the R249S mutation. Further large-scale, multi-center studies are necessary to validate our findings and establish the specific associations of these regions with the TP53 mutation at codon 249 in HCC involving AFB1 exposure and HBV infection.

## Availability of supporting data

All data generated or analyzed during this study are included in this published article and the supplementary information files.

## Acknowledgements

This work was supported, in part, by the National Nature Science Foundation of China (Grant No. 81802874,81560535). Natural Science Foundation of the Guangxi Province of China (Grant No. 2018 GXNSFBA138013). Guangxi Key R&D Program (GKEAB18221019). Key laboratory of High-Incidence-Tumor Prevention & Treatment (Guangxi Medical University), Ministry of Education (GKE2018-01).

## Authors’ Contributions

Tao Peng and Chuangye Han designed the study. Chuangye Han, Tingdong Yu and Wei Qin analyzed the data, interpreted the results and wrote the article. Tao Peng edited the article. Guangzhi Zhu, Hao Su, Xiwen Liao, Jianlu Huang, Zhengtao Liu, Long Yu, Xiaoguang Liu, Sicong Lu, Zhiwei Chen, Lequn Li, Zhen Liu, Chengkun Yang and Xiangkun Wang collected samples and clinical data. Ketuan Huang contributed to data analysis. Zengnan Mo contributed to genotyping. Tingdong Yu and Wei Qin conducted to qRT-PCR experiments. All authors discussed the results, and approved the final version of the manuscript.

## Abbreviations

HCC: Hepatocellular carcinoma
HBV: Hepatitis B virus
AFB1: Aflatoxin B1
GWAS: Genome-wide association study
TACE: Transcatheter hepatic arterial chemoembolization
BCLC: Barcelona Clinic Liver Cancer
PVTT: portal vein tumor thrombus

